# Cornelia-de Lange syndrome-associated mutations cause a DNA damage signalling and repair defect

**DOI:** 10.1101/632992

**Authors:** Gabrielle Olley, Madapura M. Pradeepa, David R. FitzPatrick, Wendy A. Bickmore, Charlene Boumendil

**Affiliations:** MRC Human Genetics Unit, Institute of Genetics and Molecular Medicine, University of Edinburgh, Crewe Road, Edinburgh EH4 2XU, UK; Blizard institute, Barts and The London School of Medicine and Dentistry, Queen Mary University of London, London, E1 2AT

## Abstract

Cornelia de Lange Syndrome is a multisystem developmental disorder typically caused by mutations in the gene encoding the cohesin loader NIPBL. The associated phenotype is generally assumed to be the consequence of aberrant transcriptional regulation. Recently, we identified a residue substitution in BRD4 associated with a Cornelia de Lange-like syndrome, that reduces BRD4 binding to acetylated histones. Here we show that, although this mutation reduces BRD4-occupancy at enhancers in mouse embryonic stem cells, it does not affect transcription. Rather it delays the cell cycle, increases DNA damage signalling, and perturbs regulation of DNA repair in mutant cells. This uncovers a new role for BRD4 in DNA repair pathway choice. Furthermore, we find evidence of a similar increase in DNA damage signalling in cells derived from NIPBL-deficient individuals, suggesting that defective DNA damage signalling and repair is also a feature of typical Cornelia de Lange Syndrome.

## Introduction

Cornelia de Lange Syndrome (CdLS) is a clinically distinctive neurodevelopmental disorder (OMIM:122470). Disease severity varies greatly and patients can suffer from a range of symptoms including: a characteristic facial appearance, upper limb abnormalities, intellectual disability and delayed growth^1^. CdLS is described as a ‘cohesinopathy’^1^ - most cases can be attributed to heterozygous loss of function mutation in *NIPBL* encoding a protein involved in loading of the cohesin complex onto chromatin^2^. Mutation in genes encoding cohesin complex proteins SMC1, SMC3 and RAD21, or HDAC8 (SMC3 deacetylase), have also been identified in CdLS-like probands^2^. However cells from CdLS patients have no obvious defects in sister chromatid cohesion^3^, and individuals with mutations in *SMC1, SMC3* and *RAD21* are often considered ‘atypical’ in terms of facial appearance and growth, and are less likely to have limb defects than those with *NIPBL* mutations^4^.

Dysregulated gene expression has been proposed to be main mechanism underlying CDLS^5,6^. Mutations in genes encoding chromatin regulators unrelated to cohesin, such as ANKRD11, KMT2A, AFF4 and the bromodomain and extra-terminal domain (BET) protein BRD4, have been reported to cause CdLS-like phenotypes^1^ suggesting that chromatin dysregulation may play a role in CdLS as well. Additionally, increased sensitivity to DNA damage has been reported in CdLS patient cells^7^, but the mechanism underlying this defect is unknown and its participation in the disease aetiology remains unclear.

Recently, we described *de novo* deletion and missense mutations in *BRD4* associated with a clinical phenotype overlapping CdLS^8^. BRD4 binds acetylated lysines residues in histones H3 and H4 through its two N-terminal bromodomain domains (BD). BRD4 localises to promoters and enhancers of active genes and is particularly enriched at super enhancers (SEs)^9,10^. BRD4 is a key regulator of transcription; through its C-terminal domain it recruits positive transcription elongation factor (P-TEFb) and the Mediator complex to promoters and enhancers, whilst its extra-terminal domain confers transcriptional activation through the recruitment of CHD4, JMJD6, and NSD3^11,12^.

The CdLS-associated BRD4 missense mutation is in the second bromodomain (BD2) (NM_058243.2:c.1289A>G, p.(Tyr430Cys), termed here as Y430C (Figure 1A), and results in decreased binding to acetylated histones^8^. To gain further insights into the mechanisms underlying CdLS, and the role of BRD4, we investigated the phenotype of mouse embryonic stem cells (mESCs) homozygous for the orthologous amino acid substitution in mouse Brd4 (actually p.Tyr431Cys but for simplicity here termed *Brd4*^*Y430C*^). Here we show that the decreased affinity for acetylated lysines results in diminished occupancy of BRD4^Y430C^ at cis regulatory elements (CREs) across the genome. However, we find no evidence of altered transcription in these cells. Instead, we report increased and more persistent DNA damage signaling and cell cycle checkpoint activation in *Brd4*^*Y430C*^ mESCs. We show increased persistent foci of the DNA damage response (DDR) protein 53BP1 upon double-strand break (DSB) induction in Brd4 mutant cells. 53BP1 is a key factor in the regulation of DNA repair pathway choice that inhibits repair by homologous recombination (HR). We also show increased foci of the downstream effectors of 53BP1, Rif1 and the Mad2l2 (Rev7) subunit of the shieldin complex in the mutant cells^13–22^ and decreased recruitment of RAD51, suggesting impaired HR repair. Further, we show that cells from CdLS patients harbouring mutations in *NIPBL* have a similar DDR phenotype, indicating there may be a previously underappreciated role for the DNA damage response in the aetiology of CdLS.

**Figure 1.**
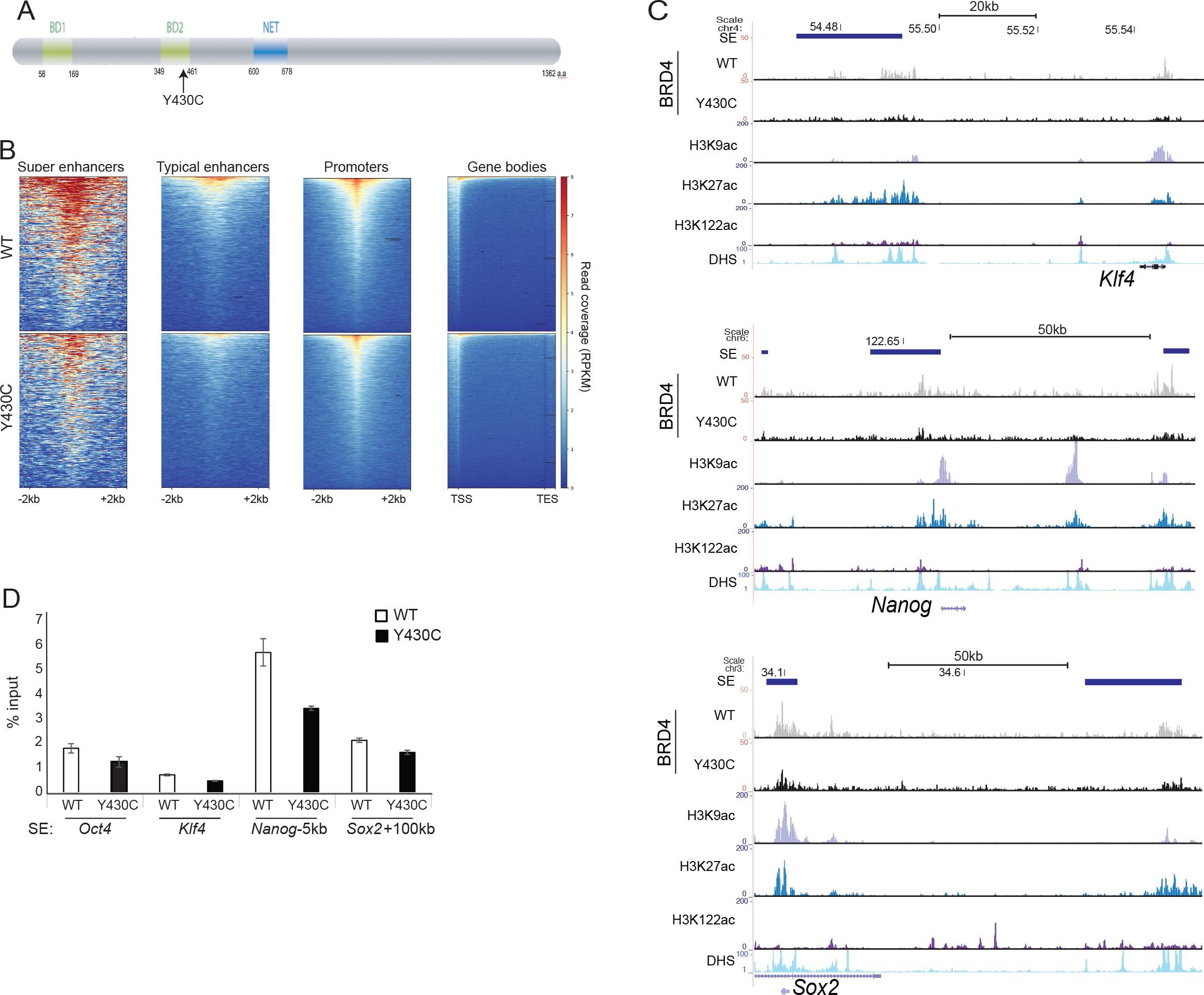
Decreased binding of BRD4 at CREs in Y430C mESCs. A) Cartoon of BRD4 showing location of the Y430C mutation in the second bromodomain (BD2). B) Heatmaps show enrichment of wild-type (WT) and Y430C BRD4 ChIP over super enhancers (SE), typical enhancers, promoters and gene bodies. C) UCSC genome browser screenshot showing reads per 10 million over the *Klf4*, extended *Nanog* and *Sox2* loci for BRD4 ChIP-seq in wild-type (WT) and BRD4^Y430C^ mESCs. Extent of SEs are shown in blue. Below are shown previously published ChIP-seq data for H3K27ac (ENCSR000CDE), H3K9ac (ENCSR000CGS), H3K122ac (GSE66023) and DNase I hypersensitivity (DHS). Genome co-ordinates (Mb) are from the mm9 assembly of the mouse genome. Biological replicate in Supplementary figure 1. D) ChIP-qPCR measuring concentration of BRD4 ChIP DNA relative to input across the SEs of *Oct4, Klf4*, *Nanog*, and *Sox2*; in WT and *BRD4*^*Y430C*^ mESCs. Data are represented as mean +/− SEM from 3 technical replicates.

## Results

### Reduced occupancy of Y430C-BRD4 at cis-regulatory elements

Our previous work suggested that the Y430C mutation abrogates BRD4 binding to acetylated histones *in vitro* and *in vivo*^8^. To determine the genome-wide effect of this loss of affinity we carried out BRD4 ChIP-seq in two independently-generated mESCs lines engineered by CRISPR-Cas9 to carry the Y430C mutation on both alleles of *Brd4*. As expected, BRD4 was enriched over CREs (SEs, typical enhancers and promoters) in both wild-type (WT) and Y430C cells (Figure 1B, Supplementary figure 1A). However, consistent with a decreased affinity for acetyl-lysines, there was a general decrease in BRD4 occupancy in Y430C cells, most striking at enhancers and super-enhancers (SE) (Figure 1C,D, Supplementary figure S1A-C). Nevertheless, Y430C binding was still sensitive to further perturbation by the BET inhibitor JQ1 (Supplementary figure 1C). In mESCs, BRD4 binding to SEs regulates the transcription of stem cell identity genes^9^. As BRD4^Y430C^ occupancy is decreased at the SEs of a number of stem cell identity genes, this suggests that there might be decreased transcription of these genes in mutant cells.

### Decreased BRD4 at CREs does not affect transcription

The use of inhibitors that competitively bind the acetyl-lysine binding pockets of BET proteins has shown that loss of BRD4 binding disrupts the expression of target genes, especially genes regulated by SEs^10^. Consistent with this, we observed decreased expression of the SE associated genes *Nanog, Myc, Klf4* and *Oct4* in WT mESCs after treatment with JQ1 (Figure 2A). However, we did not observe any decrease in levels of *Klf4*, *Nanog* and *Oct4* mRNAs in Y430C cells by RT-qPCR (Figure 2B).

**Figure 2.**
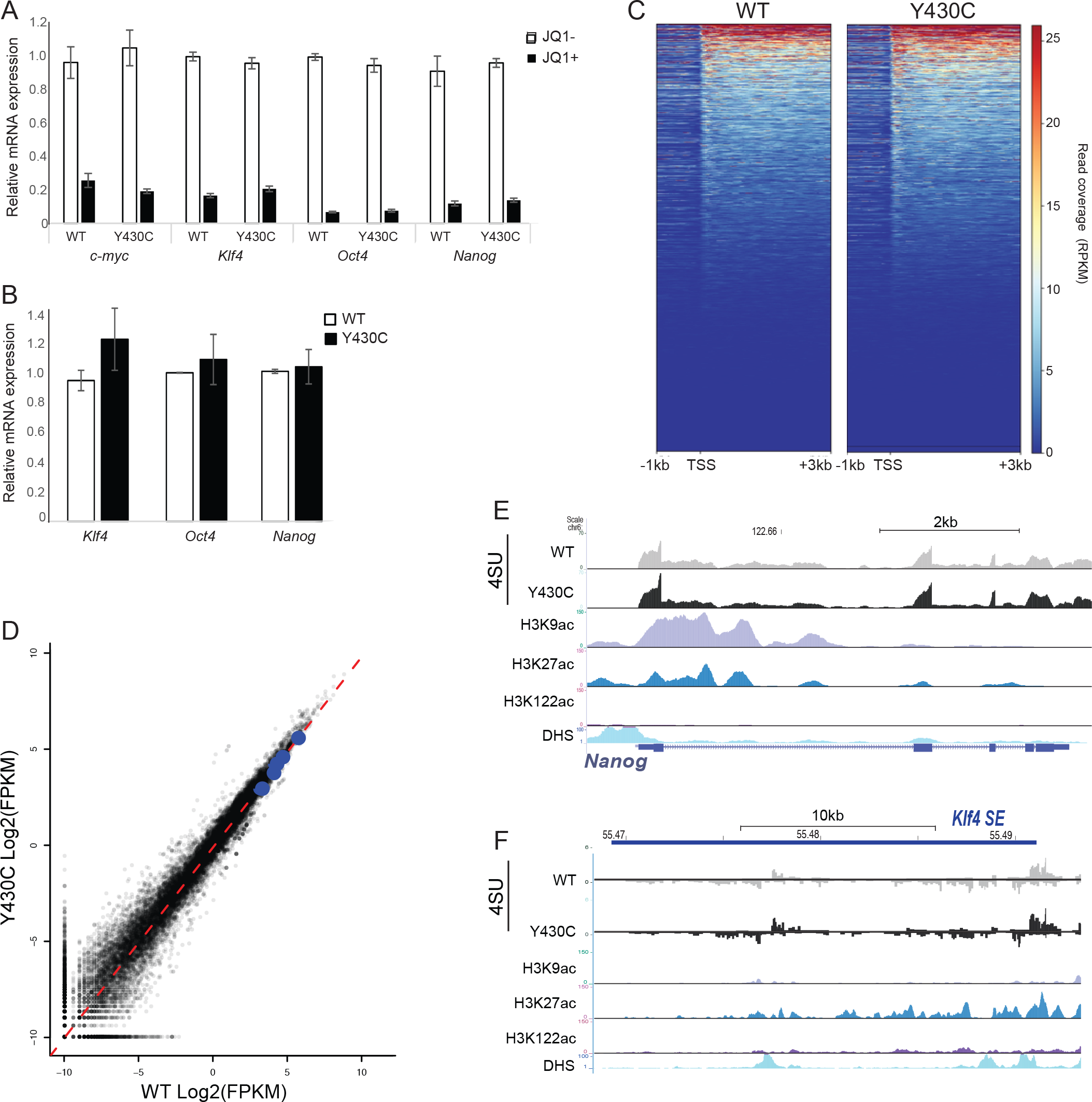
Similar transcription in WT and Y430C mESCs. A) RT-qPCR measuring mRNA of *c*-*myc, Klf4, Oct4* and *Nanog* in mESCs after treatment with 300 nM JQ1+, relative to that in untreated cells (JQ1−). Data are represented as mean +/− SEM from 3 technical replicates. B) RT-qPCR measuring mRNA for *Klf4, Oct4* and *Nanog* in WT and *BRD4*^*Y430C*^ mESCs. mRNA concentration is shown relative to WT set at 1. Data are represented as mean +/− SEM from 3 biological replicates. C) Heatmaps show enrichment of 4sU-seq in WT and *BRD4*^*Y430C*^ cells over transcribed regions (−1kb, TSS and +3kb) (mm9_refseq). D) Scatter plot of the 4SU-seq data in WT and Y430C cells, highlighting pluripotency genes in blue (*Nanog, Sox2, Klf4, Esrrb, Pou5f1*). Red dashed line shows best fitted line. Pearson correlation coefficient=0.98. E and F) UCSC browser screenshot showing 4SU-seq reads per 10 million over (E) the Nanog locus and (F) the Klf super-enhancer in WT and *BRD4*^*Y430C*^ mESCs and ChIP-seq tracks for various histone modifications and DNase I hypersensitivity in WT cells. Genome co-ordinates (Mb) are from the mm9 assembly of the mouse genome. Data from a biological replicate in Supplementary figure 2.

To determine whether mRNA stability was masking an effect on transcription per se, we performed 4-thiouridine sequencing (4SU-seq) to assay nascent transcription. Transcription was surprisingly similar between WT and Y430C mESCs (Pearson correlation coefficient=0.98) (Figure 2C,D and Supplementary figure 2A). In particular, decreased BRD4 binding at SEs did not lead to transcriptional changes at stem cell identity genes (Figure 2C-E, Supplementary figure 2B), or of eRNAs at the SEs themselves (Figure 2F, Supplementary figure 2C). Due to normalization, these experiments could not rule out that transcription is not globally decreased in the mutant cells. We therefore performed a spike-in RNA-seq experiment, using RNA from *Drosophila* cells for normalization. Again, we did not observe any major transcriptional differences between WT and Y430C cells (Supplementary figure 2D&E). We conclude that the decreased occupancy of BRD4^Y430C^ at CREs in mESCs is not sufficient to affect the transcription of associated genes. This result is surprising, given BRD4’s well documented roles in transcriptional regulation.

### Y430C-BRD4 mESCs have a delayed cell cycle and increased cell cycle checkpoint activation

We noted that *BRD4*^*Y430C*^ mESCs grew slower, and showed an accumulation of cells in G2/M (33.7%), compared to their WT counterparts (27.8%) (Figure 3A, B, Supplementary Figure 3A). This observation, together with the recently reported roles for BRD4 in the DDR and DNA repair^23–26^ led us to investigate potential DDR defects in mutant cells.

**Figure 3.**
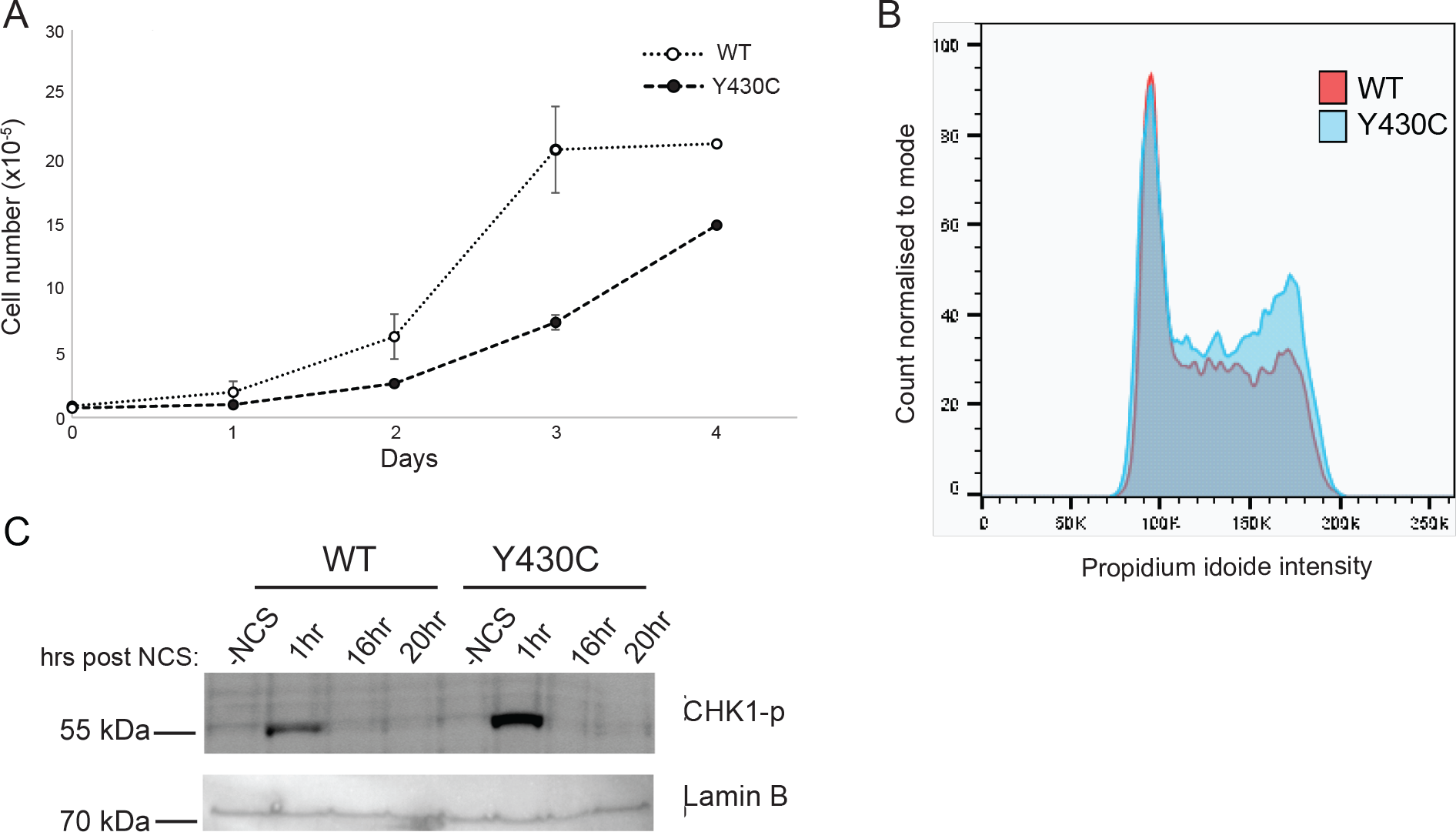
Increased G2/M checkpoint activation in Y430C mESCs. A) Graph shows average number of WT and *BRD4*^*Y430C*^ cells per well at 1, 2, 3 and 4 days post seeding. Data are represented as mean +/− SEM from 3 technical replicates. B) Overlaid graphs show WT and *BRD4*^*Y430C*^ cell cycle profiles, as determined by flow cytometry. Graphs illustrate the cell count, which correlates to propidium iodide intensity. Biological replicate in Supplementary Figure 3A. C) Immunoblot using antibodies against CHK1-p and Lamin B after treatment of WT and *BRD4*^*Y430C*^ mESCs with NCS and for various times (hrs) of recovery.

The DDR allows coordination between DNA repair and cell cycle progression. Recognition of DNA damage by sensor proteins initiates a cascade that results in the phosphorylation and activation of the checkpoint kinases CHK1 and CHK2, delaying or blocking cell cycle progression^27^. CHK1 is the main kinase required for delay at G2/M^27^. To determine whether the altered cell cycle in *BRD4*^*Y430C*^ cells is associated with increased activation of the G2/M checkpoint, we analysed CHK1 phosphorylation (CHK1-P) after treatment with neocarzinostatin (NCS), a radiomimetic drug which induces mainly DSBs. CHK1-P is increased in both WT and *BRD4*^*Y430C*^ mESCs cell lines 1hr post NCS treatment, which is resolved by 16hrs. However, the levels of CHK1-P are higher in *BRD4*^*Y430C*^ mESCs, suggesting an increased checkpoint activation (Figure 3C). There is a similar increase in CHK1-P at intermediate (2, 4, 6 and 10hr post NCS) time points discounting the possibility that checkpoint activation simply occurs faster in the mutant cells (Supplementary figure 3B).

These results suggest a defect in DNA repair or signaling caused by BRD4^Y430C^. BRD4 has been shown to be directly involved in DNA repair through the transcriptional regulation of DNA repair proteins^24,25,28^. However, 4SU-seq showed that transcription of genes encoding DNA repair proteins was unaffected in *BRD4*^*Y430C*^ mESCs (Supplementary figure 3C, D).

### Y430C-BRD4 mESCs have increased DDR signalling

BRD4 restricts the DDR and depletion of BRD4 isoform B leads to increased DDR signalling^23^. We therefore tested whether BRD4^Y430C^ affects DNA damage signalling. mESCs have constitutively high levels of yH2AX, even in the absence of a DNA damaging stimulus^29^. We therefore used 53BP1 as a marker of DDR. 53BP1 is recruited to DSBs, spreads to form microscopically visible foci, and acts as a scaffold for the recruitment of further DSB response proteins, to regulate the choice of DNA repair pathway and to promote cell cycle checkpoint signalling^30^.

Immunofluorescence showed formation of multiple 53BP1 foci, representing DNA damage sites upon DSB induction (1h after NCS treatment). These foci are only present at low levels prior to NCS treatment and decrease in number at 16 and 20h post treatment, as cells repair the damage (Figure 4A). Supporting the hypothesis that the Y430C mutation impairs the role of BRD4 role in DDR restriction, we observed that 53BP1 foci are larger in *BRD4*^*Y430C*^ mESCs than in WT (Figure 4A&B). In addition, whilst the number of 53BP1 foci in WT cells returns to pre-treatment levels at 16 and 20h time points, the number of 53BP1 foci in *BRD4*^*Y430C*^ cells remains higher (Figure 4A&C, Supplementary figure 4A&B), suggesting that DNA repair itself could be impaired.

**Figure 4.**
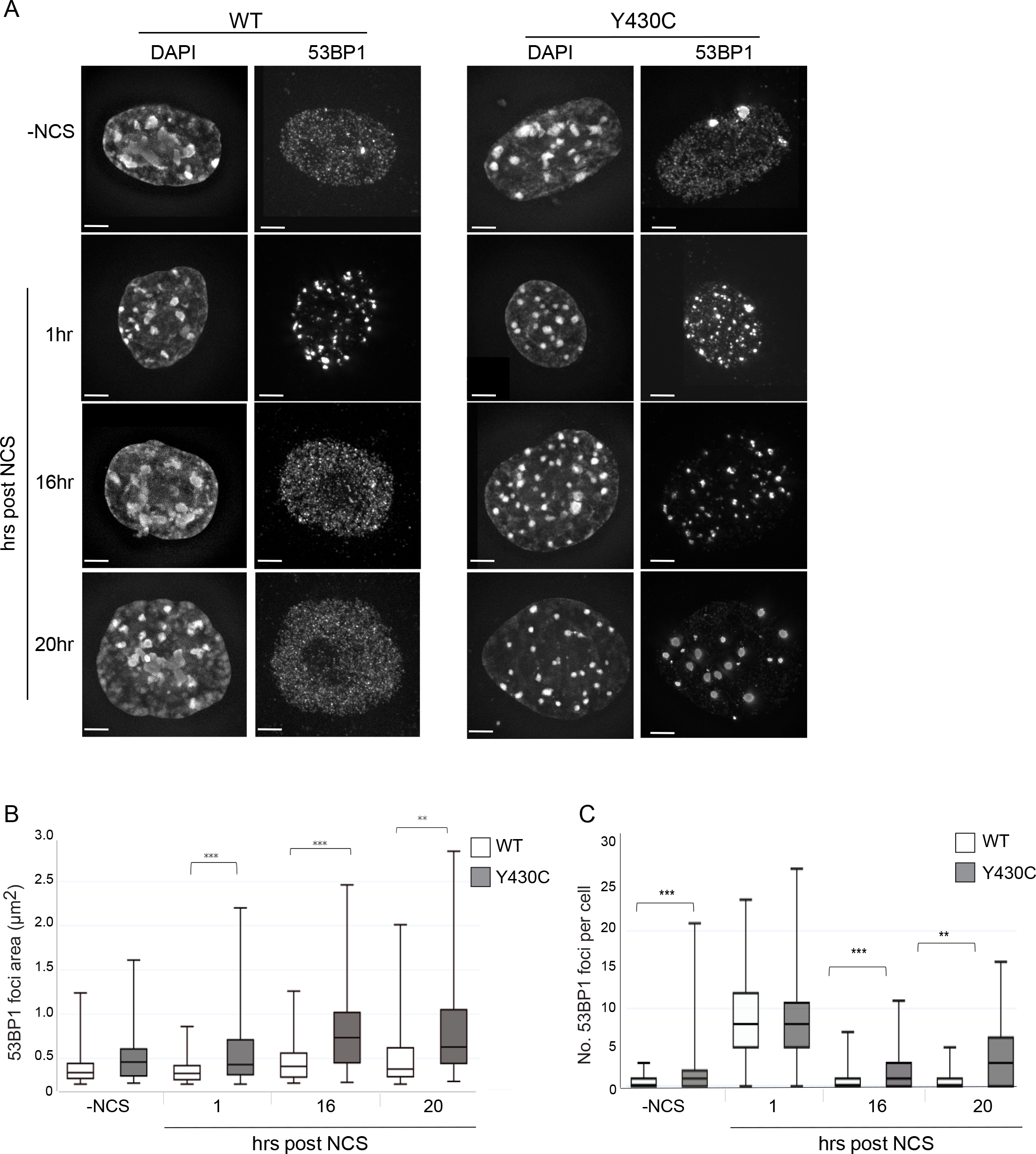
Increased size and number of 53BP1 foci after DSB induction in Y430C mESCs. A)Immunofluorescence for 53BP1 in the DAPI-stained nuclei of wild-type and *BRD4*^*Y430C*^ mESCs upon treatment with NCS and after recovery periods up to 20 hrs. B&C) Box-plots show area (μm^2^) and number of 53BP1 foci per cell, respectively, in WT and *BRD4*^*Y430C*^ cells after treatment with NCS in one representative experiment. Horizontal lines within boxes show medians, boxes are inter-quartile ranges and whiskers are range. P-values were calculated with Mann-Whitney U test. * < 0.05, ** < 0.01, *** < 0.001.

### Defective DSB repair in Y430C-BRD4 cells

For the most part, DSBs are repaired by either non-homologous end-joining (NHEJ) or HR^31^. Use of the appropriate pathway is important for faithful repair and is determined by antagonistic recruitment of 53BP1 and BRCA1^30^. 53BP1 inhibits DSB end resection, the initial step of HR, thereby promoting NHEJ and inhibiting HR. Downstream effectors of 53BP1 in the regulation of resection include RIF1^19–22^ and the recently identified shieldin complex (SHLD1, SHLD2, SHLD3 and MAD2L2)^13–18^. If timely repair does not occur by NHEJ, BRCA1 promotes the release of RIF1, leading to end-resection and HR.

As *BRD4*^*Y430C*^ mESCs show increased numbers and size of 53BP1 foci compared to WT cells, we reasoned that there may also be increased recruitment of the downstream effectors of 53BP1 such as RIF1 and MAD2L2. Indeed, we observed an increased number of RIF1 (Figure 5A&B, Supplementary figure 4C&D) and MAD2L2 (Figure 5C&D, Supplementary figure 4E&F) foci in *BRD4*^*Y430C*^ compared to WT cells at all time-points, similar to the 53BP1. Conversely, we observed a significant decrease in the number of foci of RAD51, a protein necessary for HR repair, in mutant cells at 1 hour post NCS (Figure 6A&B, Supplementary figure 4G&H), strongly suggesting a repression of HR. Given the role of the shieldin complex in protecting DSB end-resection, we propose that the Y430C BRD4 mutation leads to an altered balance between NHEJ and HR, consistent with the synthetic lethality observed between BRD4 and PARP inhibitors^25,28^.

**Figure 5.**
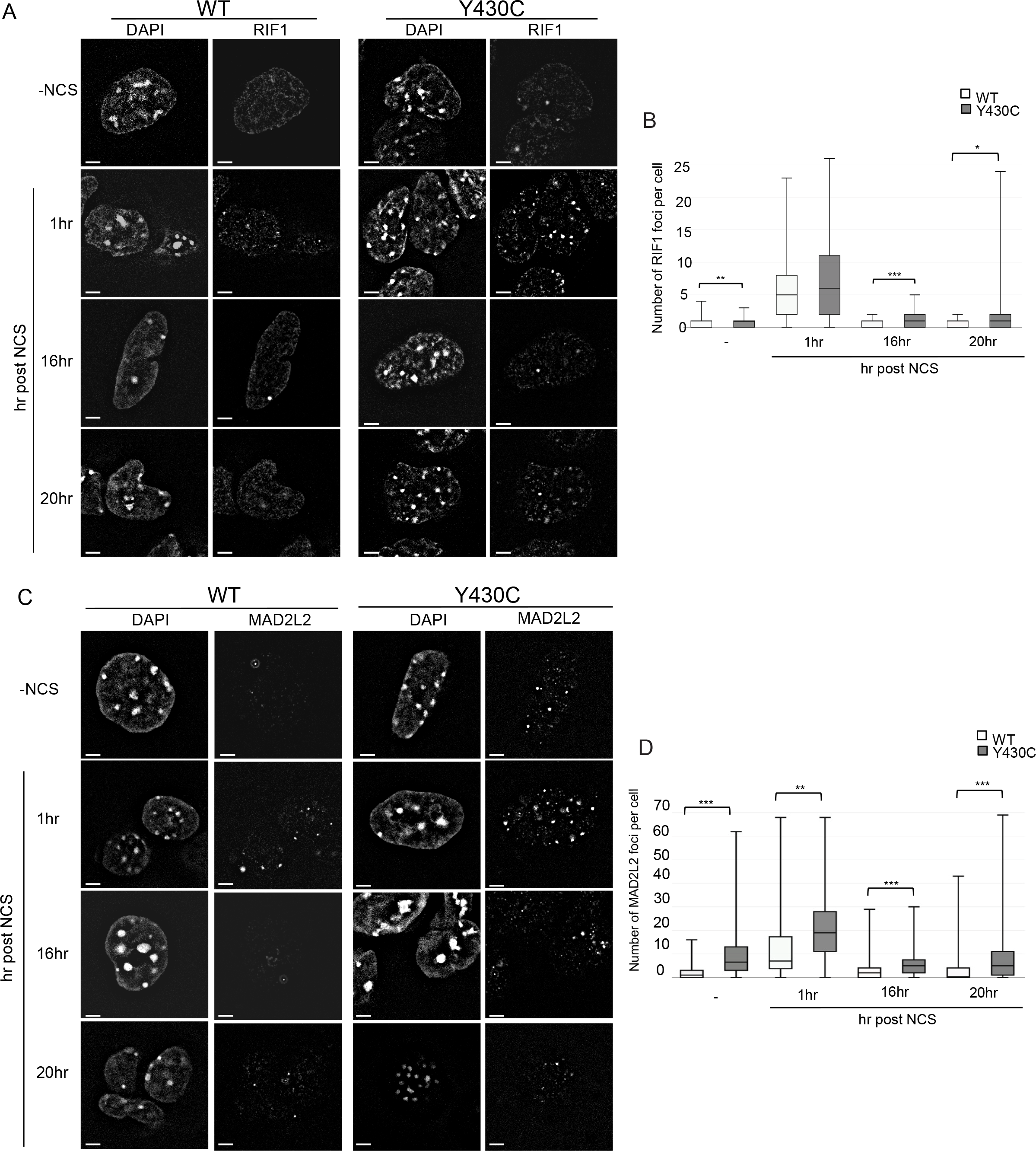
Increased RIF1 and MAD2L2 foci after DSB induction in Y430C mESCs. A) Representative images of wild-type and Y430C mESCs upon RIF1 immunofluorescence and DAPI staining after treatment with NCS. B) Box-plot shows number of RIF1 foci per cell, respectively, in WT and Y430C cells after treatment with NCS in one representative experiment. Horizontal lines within boxes show medians, boxes are inter-quartile ranges and whiskers are range. P-values were calculated with Mann-Whitney U test. * < 0.05, ** < 0.01, *** < 0.001. C) Representative images of wild-type and Y430C mESCs upon MAD2L2 immunofluorescence and DAPI staining after treatment with NCS. D) Box-plot shows number of MAD2L2 foci per cell, respectively, in WT and Y430C cells after treatment with NCS in one representative experiment. Horizontal lines within boxes show medians, boxes are inter-quartile ranges and whiskers are range. P-values were calculated with Mann-Whitney U test. * < 0.05, ** < 0.01, *** < 0.001.

**Figure 6.**
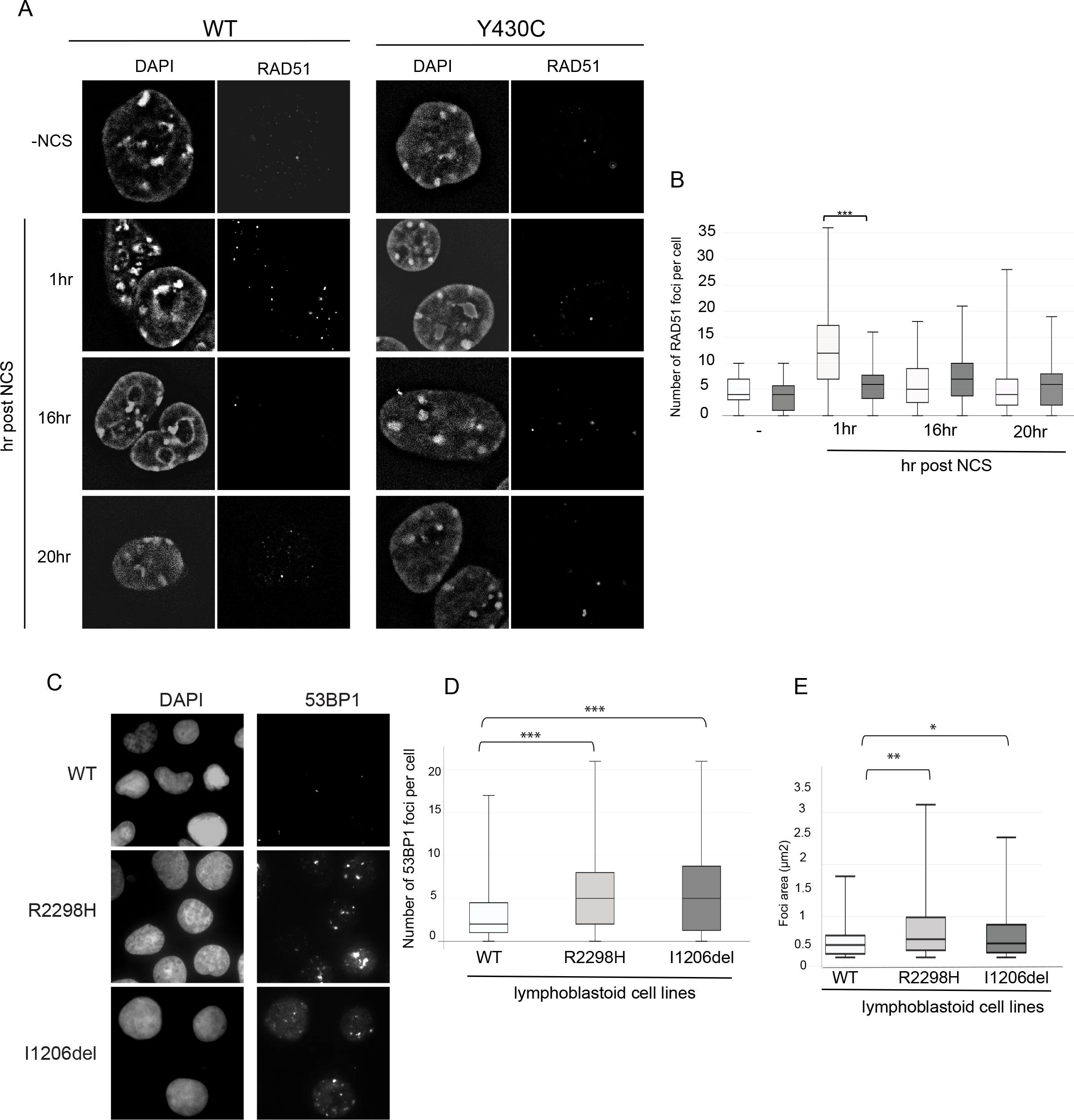
Evidence for DNA repair defects in CDLS. A) Representative images of wild-type and Y430C mESCs upon RAD51 immunofluorescence and DAPI staining after treatment with NCS. B) Box-plot shows number of RAD51 foci per cell, respectively, in WT and Y430C cells after treatment with NCS in one representative experiment. Horizontal lines within boxes show medians, boxes are inter-quartile ranges and whiskers are range. P-values were calculated with Mann-Whitney U test. * < 0.05, ** < 0.01, *** < 0.001. C) Representative images of wild-type, R2298 and I1206del LCLs upon 53BP1 and DAPI immunofluorescence. D&E) Box-plots show number of 53BP1 foci per cell and area of 53BP1 foci (μm^2^), respectively, in WT, R2298H and I1206del LCLs in one representative experiment. Horizontal lines within boxes show medians, boxes are inter-quartile ranges and whiskers are range. P-values were calculated with Mann-Whitney U test. * < 0.05, ** < 0.01, *** < 0.001.

### Increased number and size of 53BP1 foci in NIPBL mutation positive lymphoblastoid cell lines

To see if the DDR defect that we have observed in the presence of the BRD4^Y430C^ would also be apparent in cells carrying other CdLS mutations, we utilised two lymphoblastoid cell lines (LCL) previously derived from CdLS patients with heterozygous mutations in NIPBL, Ile1206del^32^ and Arg2298His^33^. These LCLs have significantly more, and larger, 53BP1 foci per cell compared to a WT LCL, in the absence of any exogenous damage (Figure 6C-E, Supplementary figure 5A&B), This suggests that increased DDR signalling could be common to CdLS cases caused by BRD4 and NIPBL mutations and that impaired DNA repair pathway choice balance may be a common mechanism underlying CdLS.

## Discussion

We previously showed that a Y430C-BRD4 mutation, and BRD4 haploinsufficiency, cause a CdLS-like syndrome^8^. Previous studies have suggested that the severe developmental phenotypes associated with CdLS are due to aberrant gene regulation. Here however, we show that BRD4^Y430C^, whilst lowering the affinity of BRD4 to acetylated lysine residues and decreasing its occupancy at enhancers and SEs, causes minimal changes in transcription, at least in mESCs, in contrast to the transcriptional changes caused by the profound loss of BRD4 binding induced by BET inhibitors. Instead, we provide evidence that BRD4^Y430C^ causes increased G2/M checkpoint activation, aberrant DDR signalling, and an altered focal accumulation of proteins that promote NHEJ and inhibit HR – 53BP1 and the shieldin complex. Conversely there is a depletion of foci containing HR proteins (Rad51), suggesting a defect in HR. Our results highlight a new role for BRD4 in the regulation of DNA repair pathway choice. Whether BRD4 mutation affects repair by HR at specific regions in the genome, or globally, remains to be investigated. For example, different levels of histone acetylation in different chromatin environments – e.g. heterochromatin vs euchromatin - upon DNA damage may recruit different amounts of BRD4^34,35^.

Could aberrant DDR and DNA repair choice account for some of the phenotypes associated with CdLS? Congenital mutation in genes involved in many different genes involved in cell cycle progression and DNA repair, are - like CdLS – generally associated with intrauterine growth retardation and short stature^36^. Similarly, microcephaly also results from mutation in genes associated with S phase progression (ATR, ATRIP1, RBBP8 - Seckl syndrome; DNA ligase IV – lig4 syndrome; XRCC4 – microcephalic primordial dwarfism^37–39^). Clinically, there is strongest phenotypic overlap between CdLS and Rubinstein-Taybi syndromes (RTS)- including arched eyebrows and other shared distinctive facial features. RTS is cause by mutations in p300 or CREBBP. These lysine acetyltransferases have recently been shown to be important for acetylating proteins involved in the DDR and DNA repair ^40^. NIPBL and cohesin are also both involved in DNA damage signalling and repair ^41^ and CdLS patient cells carrying *NIPBL* mutations display an increased DNA damage sensitivity^7^. Furthermore, we show an increase in 53BP1 foci number and size, similar to that seen in *BRD4*^*Y430C*^ mESCs, in *NIPBL* haploinsufficient LCLs. Even though we cannot discount that *BRD4* mutation in CdLS cases – Y430C, or heterozygous deletions, cause aberrant transcriptional regulation in cell types other than ESCs, our results suggest that dysregulation of DDR and repair may contribute to the aetiology of CdLS.

## Acknowledgments

We thank the individuals with CdLS and their families for generously donating their samples and genetic information and for consenting to their use in research studies. We thank Tom Strachan (Newcastle) and Matt Deardorff and Ian Krantz (Children’s Hospital of Philadelphia) for their gifts of CdLS LCLs. We thank the Institute of Genetics and Molecular Medicine Advanced Imaging Resource for assistance with imaging and Ilya Fliamer for his help with deposition of data on GEO. We thank Andrew Jackson (IGMM, Edinburgh) for feedback on the manuscript.

G.O was supported by a PhD studentship from the Medical Research Council (MRC). M.M.P. is supported by is supported by the Wellcome Trust Research seed award and funding from the School of Medicine and Dentistry, QMUL. D.R.F is supported by MRC University Unit grant (MC_UU_00007/3) and by the Simons Initiative for the Developing Brain. W.A.B is supported by MRC University Unit grant (MC_UU_00007/2). C.B was supported by a H2020 Marie-Curie Individual Fellowship (655350- NPCChr) and a Bettencourt-Schueller foundation prize for young researchers.

## Authors contributions

W.A.B, P.M.M and C.B conceived and designed the experiments with input from D.R.F. G.O conducted most of the experiments with help from P.M.M for ChIPseq and RNAseq experiments. C.B performed immunostainings and analysis of RIF1 and MAD2L2. G.O, W.A.B and C.B wrote the manuscript with input from all authors.

## Methods

### KEY RESOURCES TABLE

| Antibodies | SOURCE | IDENTIFIER |
| --- | --- | --- |
| BRD4 | Bethyl | Cat# A301-985A-M |
| 53BP1 | Novus | Cat# NB100-304 |
| Normal Rabbit IgG | Santa Cruz | Cat# sc-2025 |
| CHK1 | Abcam | Cat# ab47574 |
| CHK1-p | Cell signaling technologies | Cat# 2348 |
| Lamin B | Santa Cruz | Cat# sc-374015 |
| MAD2L2 | Abcam | Cat# ab180579 |
| RIF1 | A kind gift from Sara Buonomo | Rabbit anti-mouse Rif1 serum 1240 <sup>42</sup> |
| RAD51 | Calbiochem | Cat# PC130 |
| Goat anti-Rabbit IgG, secondary, Alexa Fluor 488 | Invitrogen | Cat# A11034 |
| Donkey anti-Rabbit IgG, secondary, Alexa Fluor 586 | Invitrogen | Cat# A10042 |

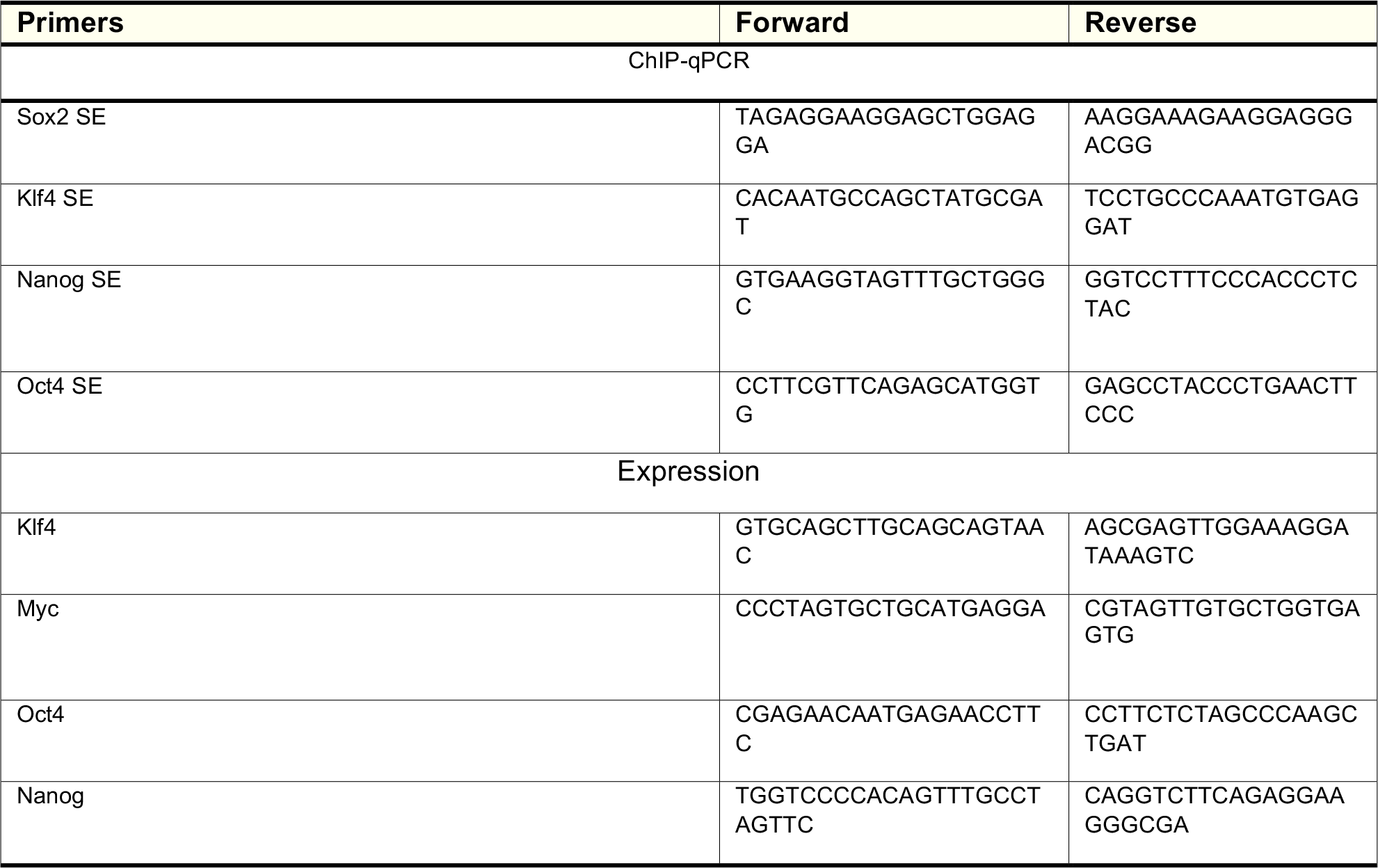

| Deposited Data |  |  |
| --- | --- | --- |
| BRD4-WT ChIP-seq | This paper | GSE130659 |
| BRD4-Y430C ChIP-seq | This paper | GSE130659 |
| WT 4sU-seq | This paper | GSE130659 |
| Y430C 4sU-seq | This paper | GSE130659 |
| WT Spike-in RNAseq | This paper | GSE130659 |
| Y430C Spike-in RNAseq | This paper | GSE130659 |
|  | This paper | GSE130659 |

| Software and Algorithms |  |  |
| --- | --- | --- |
| Bowtie2 | Langmead and Salzberg, 2012 | http://bowtie-bio.sourceforge.net/bowtie2/index.shtml |
| MACS2 |  | https://github.com/taoliu/MACS |
| FACSDiva software | BD Bioscience |  |
| TopHat | Trapnell *et al.*, 2012 | https://ccb.jhu.edu/software/tophat/index.shtml |
| Cufflinks | Trapnell *et al.*, 2012 | http://cole-trapnell-lab.github.io/cufflinks/ |
| DeepTools2 | Ramirez *et al.*, 2016 | http://deeptools.readthedocs.io/en/latest/index.html |
| SAMtools | Li *et al.*, 2009 | http://samtools.sourceforge.net/ |

### Cell culture

Y430C-BRD4 mutant and corresponding wild-type mouse embryonic stem cells (mESCs) were generated by CRISPR Cas9 genome editing in 46C mESCs as described previously ^8^. NIPBL I1206del and R2298H lymphoblastoid cell lines (LCLs) were obtained from patients^32,33^. mESCs were cultured in GMEM medium (GIBCO; 11710035) supplemented with 10% Fetal Calf Serum (FCS), 5% penicillin-streptomycin, 1 mM sodium pyruvate (GIBCO; 11360070), 1X non-essential amino acids (GIBCO; 11140050), 50 μM 2-Mercaptoethanol (GIBCO; 31350010), 2 mM L-glutamine and 500U/ml Leukaemia Inhibitory Factor (in house). Lymphoblastoid cell lines (LCLs) were grown in RPMI 1640 medium (GIBCO; 11875093) supplemented with 15% FCS and 2 mM L-glutamine. All cells were grown at 37°C in a 5% CO_2_ humidified atmosphere.

### ChIP-qPCR

Cells were harvested by trypsinising and fixed with 1% formaldehyde (Thermo Fisher; 28906) in media (25°C, 10 min). This reaction was quenched with 0.125 M glycine for 5 min. ChIP-qPCR was performed as described previously^8^ (see table for antibodies). DNA was purified using the QIAquick PCR Purification kit (Qiagen, 28104). Input samples were diluted to 1%, and all samples diluted a further 10-fold, in ddH20. SYBR-green based qPCR reactions were performed in a final volume of 20 μl containing diluted ChIP DNA, SYBR select master mix (ThermoFisher Scientific; 4472908) and 0.25 μM/L of each primer (see table). Concentration of IPs are relative to 1% input.

### ChIP-seq

ChIP was carried out as above. After purification, DNA was eluted in 20 μl and libraries were prepared for ChIP and input samples as previously described^43^. Samples were sequenced at BGI (Hong Kong; 50-bp single-end reads) using the HiSeq 4000 system (Illumina). Fastq files were quality controlled using FastQC and mapped to the mm9 genome using Bowtie2 (parameters: default). Sam files were converted to bam files and sorted using SamTools. Homer was used to make tagdirectories (makeTagDirectory, parameters: -unique, fragLength 150) and bedgraphs (makeUCSCfile, parameters: default). For visualisation of BRD4 data, bedgraphs were uploaded to the genome browser UCSC. Peak calling was carried out using MACS2; Duplicates were filtered (filterdup, parameters:--keep-dup=1), peaks called (callpeaks, parameters: -B --nomodel -p 1e-5) and differential peaks were found (bdgdiff, parameters: -g 60 -l 250).

deepTools2 was used to make heatmaps; score files were made across specific genomic regions (computeMatrix, parameters: scale-regions scale regions -b 500 -a 500 -bs 50 -bl mm9 blacklist) and these were used to plot heatmaps (plotHeatmap, parameters: --colormap RdBluYl reverse).

### JQ1 treatment

1 mM BRD4 inhibitor JQ1+, or its inactive form JQ1− (Merck; 500586) (diluted in DMSO), were added to mESC media at a final concentration of 300 nM. JQ1+/-. WT and Y430C mESCs were incubated at 37°C with JQ1+/− supplemented media for 48 hrs. Total RNA was extracted from cells using the RNeasy Plus Mini Kit (Qiagen; 74134) and 1 μg RNA was used for cDNA synthesis with SuperScript II Reverse Transcriptase (ThermoFisher Scientific; 18064-014) as per manufacturer’s instructions. cDNA was diluted 1:500 for qPCR analysis. qPCR reactions were performed as above (see table for primers). Concentration of JQ1+ cDNA was calculated relative to JQ1− (arbitrarily set to 1).

### RT-PCR

RNA was extracted from cells using the RNeasy Mini Kit (Qiagen; 74104) using spin technology, with an additional on-column DNA digestion using the RNase-Free DNase Set (Qiagen; 79254). cDNA was synthesised from 1 μg RNA using SuperScript II Reverse Transcriptase (ThermoFisher Scientific; 18064-014) as per manufacturer’s instructions. cDNA was diluted 1 in 25 for qPCR analysis. SYBR-green based qPCR reactions were performed in a final volume of 20 μl containing diluted cDNA, SYBR select master mix (ThermoFisher Scientific; 4472908) and 0.5 μM/l of region specific intron-spanning primer pairs.

### 4sU-seq

4sU RNA was generated and isolated as described previously ^44^, with the following changes: cells were incubated at 37°C with 4sU-supplemented medium for 20 min. The reaction was incubated with Biotin-HPDP with rotation for 1.5 hours at RT. For recovery of biotinylated 4sU-RNA, 1 μl of streptavidin beads was added per μg of RNA. Columns were washed using 900 μl washing buffer and RNA was eluted by 2 sequential additions of 100 μl Elution Buffer (100 mM DTT) to the column and eluates combined. RNA was further purified using the RNAeasy MinElute Clean-up kit (Qiagen; 74204) according to the manufacturer’s guidelines, eluting in 20 μl water. 1 μl of 4sU-labeled RNA was quality-checked by running on a 2100 Bioanalyzer Instrument (Agilent).

To make 4sU sequencing libraries, 4sU labelled RNA was first depleted of rRNA using the Low Input Ribominus Eukaryotic System V2 (ThermoFisher Scientific; A15027) as per the manufacturer’s instructions. 600 ng of 4sU labelled RNA was used as input, and eluted in 5 μl RNase free water. All of the resulting rRNA free RNA was used to prepare 4sU sequencing libraries, using NEBnext Ultra Directional RNA library prep kit of Illumina (NEB; E7420). RNA fragmentation was carried out at 94°C for 15 min, as suggested for intact RNA. Libraries were indexed with Multiplex Oligos for Illumina^®^ (Index Primers Set 1) (NEBnext; E7335) and amplified by PCR for 13 cycles. Library concentration and correct size distribution was confirmed on the Agilent 2100 Bioanalyser with the DNA HS Kit. Libraries were sequenced at BGI (Hong Kong; 100-base paired-end reads) using the HiSeq 4000 system (Illumina).

Fastq files were quality controlled using FastQC and mapped to the mm9 genome using tophat (parameters: --library-type fr-firststrand -r 200). Homer was used to make tagdirectories (makeTagDirectory, parameters: -unique -sspe -flip -fragLength 150), and to make bedgraphs for visualisation on UCSC (makeUCSCfile, parameters: -strand separate -style rnaseq). Cufflinks was used for peak calling; transcripts were assembled for individual experiments (cufflinks, parameters: –m 200 –library-type fr) and both replicates of WT and Y430C were combined to form one assembly (cuffmerge, parameters: default). Differentially expressed peaks were determined from this assembly using cuffdiff (Cuffdiff. Parameters: default).

Heatmaps were generated as above.

### Spike-in RNA-seq

S2 cells were cultured in Schneider’s Drosophila Medium (Invitrogen; 11720-034), supplemented with 10% heat-inactivated FCS and 5% penicillin-streptomycin. Cells were passaged once they reached a density of ~2×107 cells/ml and seeded at a density of ~4×106. Cells were grown at 28°C in a 5% CO2 humidified atmosphere. Cells were frozen at a density of ~1×107 cells/ml in 45% conditioned Schneider’s Drosophila Medium media (containing 10% FCS), 45% fresh Schneider’s Drosophila Medium supplemented with 10% FCS, and 10% DMSO, and stored in liquid nitrogen.

mESCs and S2 cells were harvested and counted. 0.2 million S2 cells were mixed with 10 million mESCs, and RNA was extracted using the RNeasy Mini Kit (Qiagen; 74104) using spin technology, with an additional on-column DNA digestion using the RNase-Free DNase Set (Qiagen; 79254). RNA was depleted of rRNA and RNA-seq libraries prepared as for the 4sU-seq.

### Growth assay

WT and Y430C mESCs were each seeded in 4 wells of a 6 well plate (1 × 10^4^ cells/well). WT and Y430C cells from 1 well were trypsinised and counted at 24, 48, 72 and 96 hrs post seeding. Counting was carried out manually using a haemocytometer. The addition of trypan blue dye allowed for the exclusion of dead cells.

### Flow cytometry

2 million mESCs were fixed in 70% ethanol (in PBS) at 4°C for 1 hr. Fixed cells were centrifuged at 2000 rpm at 4°C for 5 min, washed twice with PBS and resuspended in 500 μl PBS. 20 μg RNase A was added and cells were incubated at 37°C for 10 min. Cells were stained with propium iodide at a final concentration of 50 μg/ml. Acquisition was carried out on a BD LSRFortessa cell analyser, collecting 25,000 events per sample. Results were analysed using BD FACSDiva 8.0.1 and gated cells were manually categorized into cell cycle stages G0/G1, S and G2/M.

### NCS treatment and CHK-1 protein western blots

Cells were incubated with mESC media supplemented with neocarzinostatin (Sigma; N9162) (NCS), to a final concentration of 25 ng/ml, for 15 min at 37°C. Cells were then washed with PBS and fresh, non-supplemented media was added. Protein was either extracted straight away, or after incubation at 37°C for varying lengths of time. Ice-cold RIPA buffer (150 mM sodium chloride; 1.0% NP-40; 0.5% sodium deoxycholate; 0.1% SDS; 50 mM Tris, pH 8.0) was added to plates (1 ml per 10^7^ cells) and cells were scraped and transferred into pre-chilled microcentrifuge tubes. Tubes were shaken at 4°C for 30 min before centrifugation at 20,000 x *g* for 15 min. Supernatant was retained and quantified. For western blot analysis, equal amounts of protein were boiled in 1X NuPage LDS buffer (ThermoFisher Scientific, NP0008) with 1X NuPage reducing agent (ThermoFisher Scientific; NP0004) for 5 min and separated on a 3-8% tris-acetate gel (ThermoFisher Scientific; EA0375BOX). Following electrophoresis, proteins were transferred to nitrocellulose membranes (ThermoFisher Scientific) and immunoblotted with primary antibodies overnight at 4°C. Membranes were washed 3 X TBST and probed with HRP-conjugated secondary antibody for 1 hr at RT. After 3 more washes in TBST, membranes were incubated with SuperSignal™ West Femto Maximum Sensitivity Substrate (Thermo Fisher Scientific; 34095) for 5 min and imaged using ImageQuant™ LAS 4000 (GE Healthcare).

### Immunofluorescence

mESCs for immunofluorescence experiments were cultured on gelatinised coverslips and LCLs were grown in suspension. LCLs were harvested and resuspended in PBS to 1.8 × 10^5^ cells/ml. 500 μl of cell suspension was added to a Shandon™ Single Cytofunnel™ (Thermo Fisher Scientific; 5991040), with a microscope slide attached. Slides were centrifuged at 800 rpm for 5 min, after which the LCLs had attached to the slide. All cells were fixed in 4% paraformaldehyde for 10 min and washed 3X 3 min in PBS. Cells were then permeabilised in 0.5% Triton in PBS for 10 min and washed 3X 3 min in PBS. Cells were blocked in 1% BSA in PBS for 30 min at RT, incubated with primary antibody diluted in 1% BSA for 1 hr at RT and washed 3X 3 min in PBS. Cells were next incubated with secondary antibody (see table) diluted in 1% BSA for 45 min at RT, washed 3X 3 min in PBS, incubated with DAPI in PBS (250 ng/ml) for 2 min, and washed 3x 3 min in PBS. Coverslips were mounted on slides in Vectashield (Vector; H1000) mounting medium for fluorescence.

All slides were viewed, and foci counted, using epifluorescence microscopes. Images were taken using confocal microscopy.

